# Cytonemes with complex geometries and composition extend into invaginations of target cells

**DOI:** 10.1101/2020.10.09.332676

**Authors:** Brent M. Wood, Valentina Baena, Hai Huang, Mark Terasaki, Thomas B. Kornberg

## Abstract

Cytonemes are specialized filopodia that mediate paracrine signaling in Drosophila and other animals. Studies using fluorescence confocal microscopy (CM) established their general paths, cell targets, and essential roles in signaling. To investigate details unresolvable by CM, we used high pressure freezing and electron microscopy to visualize cytoneme structures, paths, contents, and contacts. We observed cytonemes previously seen by CM in the Drosophila wing imaginal disc system, including in disc, tracheal air sac primordium (ASP), and myoblast cytonemes, and identified cytonemes extending into invaginations of target cells and cytonemes connecting ASP cells and connecting myoblasts. Diameters of cytoneme shafts vary between wide (206 ± 51.8 nm) and thin (55.9 ±16.2 nm) segments at regular intervals. Actin, ribosomes, and membranous compartments are present throughout; rough endoplasmic reticulum and mitochondria are in wider proximal sections. These results reveal novel structural features of filopodia and provide a basis for understanding cytoneme cell biology and function.

**Summary:** Cytoneme signaling filopodia of Drosophila cells have regions defined by oscillating diameters, and contents that include ER, mitochondria and ribosomes, and form contacts along invaginations of target cell membranes.

## Introduction

Cytonemes are thin, actin-based filopodia that distribute morphogen signaling proteins in developing tissues (Kornberg, 2014a). They directly link signal producing and signal receiving cells in the Drosophila wing imaginal disc (Chen et al., 2017; González-Méndez et al., 2017), abdominal histoblast nest (Bischoff et al., 2013), and gonadal stem cell niche (Rojas-Ríos et al., 2012), as well as the zebrafish neural plate (Stanganello et al., 2015; Mattes et al., 2018), chick limb bud (Sanders et al., 2013), and human embryonic stem cells (Junyent et al., 2020). Signaling by morphogen proteins such as Hedgehog (Hh), Decapentaplegic (Dpp, a Bone Morphogenic Protein (BMP) family member), Wingless (Wg/Wnt), and Branchless (Bnl, a Fibroblast growth factor family member) is dependent on cytonemes and the contacts they make with target cells (Kornberg, 2017). Despite the prevalence and critical roles of cytonemes, their ephemeral nature and wispy structure has limited their physical characterization.

The contacts cytonemes make with target cells share both structural and functional attributes with neuronal synapses. The contacts are < 40 nm, close enough for complementary fragments of GFP at the surface of presynaptic and postsynaptic cells to reconstitute (Roy et al., 2014; Huang et al., 2019). The contacts are constituted with proteins that function at neuronal synapses, are excitatory, and glutamatergic (Huang et al., 2019; Roy et al., 2014). And they are sites where signals exchange between transmitting and recipient cells, and are essential for signaling (Mattes et al., 2018; Roy et al., 2014; González-Méndez et al., 2017). Further parallels include the growth cone filopodia of extending axons, which involve signaling proteins and receptors and involve signaling molecules such as Wnts, Hhs, and BMPs (Zou and Lyuksyutova, 2007). These parallels between neuronal synapses and cytoneme contacts encouraged us to ask whether ultrastructural analysis might reveal further similarities.

Electron microscopy (EM) studies of several types of membrane extensions have been reported. These include studies of thin, actin-based extensions such as transzonal projections (TZPs) (Baena and Terasaki, 2019), tunneling nanotubes (TNTs) (Rustom et al., 2004; Sartori-Rupp et al., 2019; Alarcon-Martinez et al., 2020), and axonal growth cone filopodia (Bastiani and Goodman, 1984). Serial transmission EM (TEM) analysis of grasshopper neuron growth cones identified filopodia inserting deep into invaginations of target cell membranes (Bastiani and Goodman, 1984). These burrowing extensions induce formation of coated pits and vesicles in the cell membranes. In mouse ovarian follicles, TZPs of granulosa cells contact the oocyte through gap junctions, and they have been proposed to facilitate adherens junction-mediated paracrine signaling to drive cumulus cell differentiation (Baena and Terasaki, 2019). Other granulosa cells not in contact with the oocyte project filopodia which end by inserting into invaginations of neighboring granulosa cells without membrane fusion.

The TNT term refers to a diverse set of cell protrusions that have been characterized almost exclusively in adherent cultured cells (Dupont et al., 2018). Like cytonemes, they are actin-rich, cytoplasmic cell-cell connections, and because they have been distinguished mostly by their lack of adherence to matrix or plastic growth substrates (Dupont et al., 2018; McCoy-Simandle et al., 2016), it is not clear if they are a type of filopodia distinct from cytonemes (Yamashita et al., 2018). EM analysis of TNTs in cultured neuronal cell lines reveals individual TNTs extending in parallel bundles held together by N-Cadherin linkages. TNTs have outward bulging sections with mitochondria and other membranous compartments, and most make open-ended contacts with target cells. Others are close-ended and burrow into invaginations of target cells (Sartori-Rupp et al., 2019). A SEM study of pericytes surrounding adult mouse retinal capillaries identified individual TNTs with gap junctions at close-ended foot-like processes (Alarcon-Martinez et al., 2020). Membrane nanotubes have also been characterized in virus-infected T cells. In this context, viral transfer to uninfected cells is mediated by membrane nanotubes and is receptor- and contact dependent (Sowinski et al., 2008).

Another type of cellular extension involved in signaling is the vertebrate primary cilium, which localizes specific receptors and signal transduction proteins (Anvarian et al., 2019; Kornberg, 2014b). Although the receptor and lipid composition of primary cilia membranes are specialized and regulated, primary cilia have no ribosomes (Rosenbaum and Child, 1967) and their protein constituents selectively enter through structures in the transition zone of the basal body. EM studies of cilia have identified their highly organized ultrastructure and functionally and structurally distinct regions (Fisch and Dupuis-Williams, 2011).

Using HPF and electron microscopy, we examined the structure of cytonemes and their contact points to determine if they share features with other types of signaling membrane extensions and synapses. Compared to chemical fixation, HPF better preserves tissue morphology (Studer et al., 1992), providing images of organelles and membranes that have smoother more regular appearance (Kaneko and Walther, 1995; Royer and Kinnamon, 1996), denser and more homogenous cytoplasmic matrices (Kaneko and Walther, 1995), and better preservation of physiological extracellular space (Korogod et al., 2015). Following HPF and freeze substitution, we used both serial scanning EM (SEM), which more easily allows for serial section collection, and TEM, which provides higher resolution to track cytonemes that extend from three different cell types, and visualized their connections to target cells. We found that cytonemes are present in greater numbers and diversity than CM had revealed and that they have distinctive shapes that are characterized by alternating thin and wide sections. Cytonemes and their contact points contained ribosomes, though there was heterogeneity in the amount. Membranous compartments of different sizes and shapes were present along the length of cytonemes, while mitochondria and rough ER were more confined to wider areas of membrane extensions that were proximal to the projecting cell. Cytonemes connecting ASP cells to each other and cytonemes connecting myoblastst to each other were identified in addition to previously studied cytoneme populations.

## Results

### ASP and myoblast cytonemes project towards multiple cell types

Cytoneme lengths, orientations, and dynamics have been studied in the wing disc and other systems using membrane-targeted fluorescent markers, and the role of cytonemes in signaling has been analyzed genetically and analyzed using fluorescently tagged ligands and receptors (Kornberg, 2014a). Bipartite GFP reconstitution experiments show that cytonemes make close contact with target cells. Yet the structure and dimensions of cytonemes and the nature of the cellular contacts have not been resolved because their thin diameter is not resolved by light microscopy and they are not well preserved by chemical fixation. To overcome these issues, we used HPF to preserve the wing disc rapidly prior to chemical fixation, and serial SEM and TEM to resolve the fine structures of cytonemes, the cells from which they project and to which they connect.

The wing disc is a flattened epithelial sac with a layer of columnar epithelial cells on one side that includes the primordia for the wing blade and the dorsal exoskeleton of the adult thorax (notum) (Fig. 1A). The notum primordium has several deep folds and its basal side is attached to the air sac primordium (ASP), myoblasts, and extracellular matrix. The ASP is a tubular extension of approximately 120 tracheal epithelial cells that extends posteriorly across the wing disc from the transverse connective tracheal branch. It is a single cell-layered epithelial tube with a thin proximal stalk and bulbous distal end. Myoblasts proliferate to cover most of the notum primordium during the third instar and some lie between the disc and ASP.

**Figure 1.**
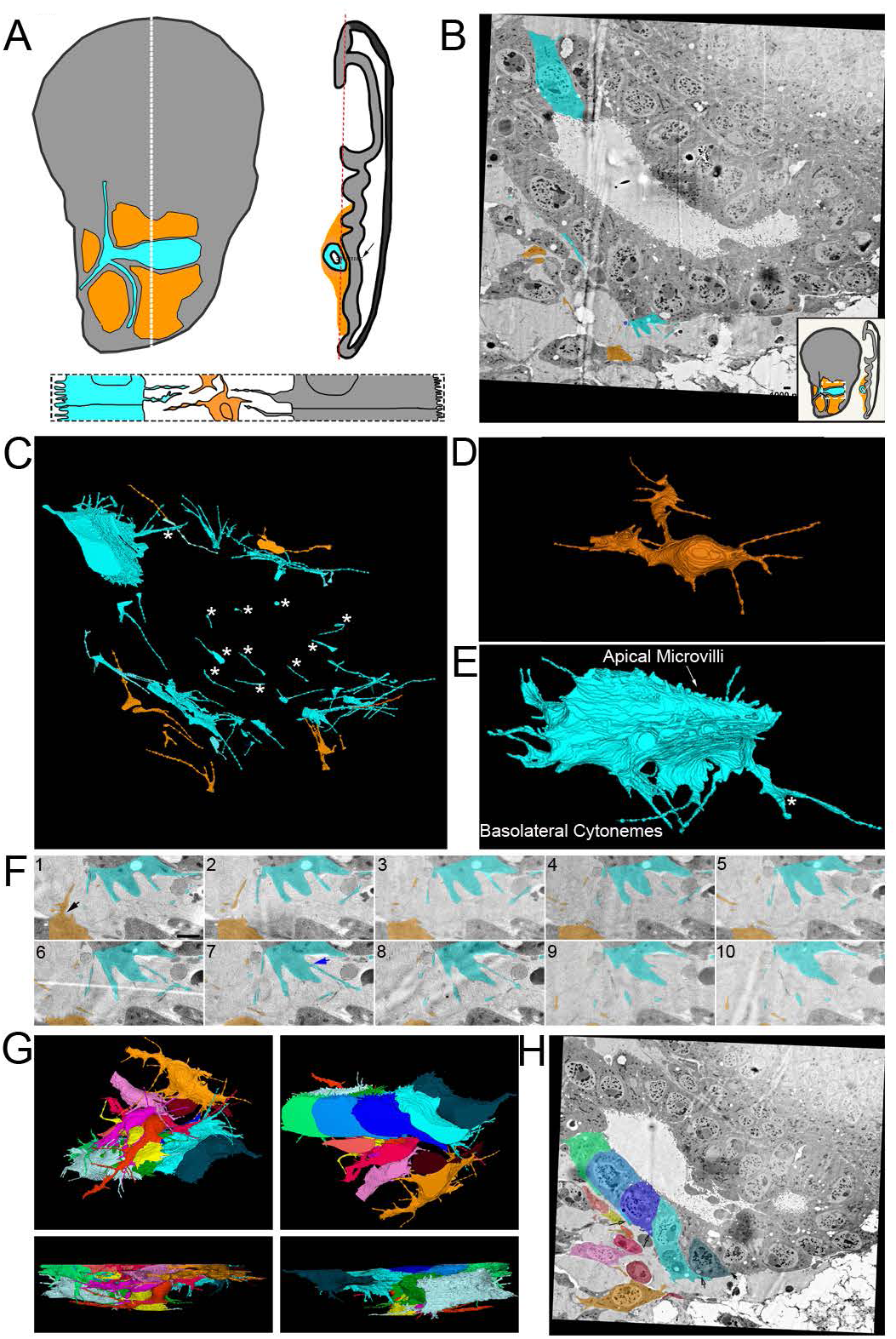
Cytonemes project from ASP epithelial cells and myoblasts. **A)** Illustration depicting the Drosophila wing imaginal disc (grey), ASP (cyan), and myoblasts (orange) from frontal (left) and sagittal (right) perspectives. The dashed lines indicate the position of the sagittal (white) and frontal (red) sections. An enlargement (bottom) of the black box in the sagittal view depicts the microvilli on the apical surfaces, and cytonemes on the basal surfaces, of both ASP and disc epithelial cells. The intervening myoblasts project cytonemes in multiple directions. **B)** One section of 172 serial SEM images showing the ASP surrounded by myoblasts. Disc epithelial cells are visible in the top right comer. The imaged region is indicated by a white box and line on the inset diagram. Each image was segmented to highlight a portion of cytoneme projecting from ASP (cyan) and myoblasts (orange). **C)** Segmentation was used to create a 3D-reconstruction of the cytoneme projecting from ASP cells (cyan) and myoblasts (orange). One reconstructed ASP cell is included in the 3D reconstruction. Asterisks indicate cytonemes projecting in the direction of cells of the same type. **D)** A reconstructed myoblast projecting cytonemes in multiple directions. **E)** A reconstructed ASP cell with microvilli on its apical surface, and cytonemes projecting from its basolateral surfaces. The asterisk indicates a cytoneme which bifurcates. **F)** An enlarged region from segmented serial SEM images showing cytonemes projecting from an ASP cell and a myoblast in the direction of the other cell type. The black arrow (section 1) indicates a cytoneme extending directly from the cell body, while the blue arrow (section 7) indicates a cytoneme extending from a thick membrane extension. Scale bars: 1 µm (B), 500 nm (F).

Cytonemes extending from the various cells of the wing disc are distinguished by the cell types they connect and by the signaling proteins they transmit. Disc cell cytonemes connect to other wing disc cells, to ASP cells, or to closely apposed myoblasts (Huang and Kornberg, 2015). The cytonemes transport Hh and Dpp between disc epithelial cells, Hh, Dpp and Bnl from disc to ASP cells, Hh and Wg between disc cells and myoblasts, and Delta between myoblasts and ASP cells.

To examine cytonemes at high resolution, we dissected wing discs gently to preserve physical associations with the ASP and myoblasts, placed the specimens in appropriate chambers, and subjected the tissues to rapid high pressure freezing. Frozen blocks were processed by freeze substitution, embedding, and sectioning, and were examined by SEM and TEM. Filopodia were observed extending from cells throughout the samples; some connected to other cells. We do not know if the free-ended extensions may be cytonemes that had disconnected from a target cell or were in the process of extending to a target, or alternatively if they may be filopodia with another function. For this paper, we refer to all filopodia in our samples as cytonemes.

We first sought to visualize and track cytonemes extending from ASP cells, myoblasts, and disc epithelial cells (Fig. 1A). To examine cytonemes extending from the ASP and myoblasts, 172 serial SEM images (Fig. 1B, Sup Fig. 1) were aligned to build a stack 10.32 microns deep. Representative cytonemes and cells were segmented and reconstructed in 3D to determine directions of projection (Fig. 1C). Whereas myoblasts are not polarized cells, and their cytonemes project from all sides (Fig. 1D, Supplemental Movie 1), the epithelial ASP cells project cytonemes from lateral and basal surfaces in the direction of myoblasts (Fig. 1E,F, Supplemental Movie 1). ASP cells also have microvilli on their apical surface (Fig 1E, Supplemental Movie 1). Some of the myoblast cytonemes project in the direction of the ASP (Fig. 1C and E). Cytonemes were also observed projecting between cells of the same type (Fig. 1C, asterisks). In some instances, a single cytoneme projects from a cell before bifurcating (Fig. 1E, asterisk). Some cytonemes project directly from the cell body (Fig. 1F, section 1 arrow), while others project from thicker membrane extensions (Fig. 1F, section 7 arrow). Most cytonemes terminate without contacting another cell.

Reconstruction of seven ASP cells and eight nearby myoblasts (Fig. 1G) revealed regions with extracellular space (Fig. 1H) as well as closely apposed surfaces. The longest cytonemes these cells project are 8.28 µm (ASP) and 12.4 µm (myoblast), although most are <7 µm. 11.1% of the cytonemes extend outside the imaged area and their complete length could not be measured. By CM, cells at the tip of the ASP have been shown to extend cytonemes up to 40 µm (Roy et al., 2014). These tip ASP cells were not present in their entirety in the serial SEM stack. The imaged area, which was approximately 58 × 58 x 10 µm, allowed us to examine cytonemes projecting between the ASP and surrounding myoblasts but precluded finding the longest cytonemes between ASP and disc cells.

Although ASP cells project thick extensions and cytonemes in many directions, a thick basal extension that tracks distally is a consistent ASP cell feature. Most ASP cells, including both cells that were and were not 3D reconstructed, have a substantial portion of their basal surface within the imaged area. Of those cells, 93% project a thick membrane extension (Fig. 1H, arrows, Supplemental Movie 1) that tracks distally along the surface of the neighboring cell. 98% of those thick extensions narrow to cytonemes, of which the majority (79%) continue to project distally. The myoblasts project cytonemes in multiple directions, both towards and away from the ASP. Using serial SEM we visualized previously identified populations of ASP and myoblast cytonemes that extend between these cell types; we also identified cytonemes that project in the direction of cells of the same type.

### Cytonemes extend from basal and lateral surfaces of disc epithelial cells

We next visualized cytonemes extending from the disc where basolateral cytonemes have been previously analyzed by CM (Callejo et al., 2011). At the basal side of the disc, cytonemes project from epithelial cells (Fig. 2A-B). These cytonemes appear similar in size and shape to those of the ASP and myoblasts. At the basal side of a fold region, a series of ten sections was aligned, segmented (Fig. 2C) and reconstructed in 3D (Fig. 2D). Cytonemes extend along the fold (i.e. aligned with the anterior/posterior axis), and there are no cytonemes perpendicular to the fold. Thicker membrane extensions are also present, and some of these membrane extensions transition from thick to thin within the stack (Fig. 2D, pink and Fig. 2E, teal).

**Figure 2.**
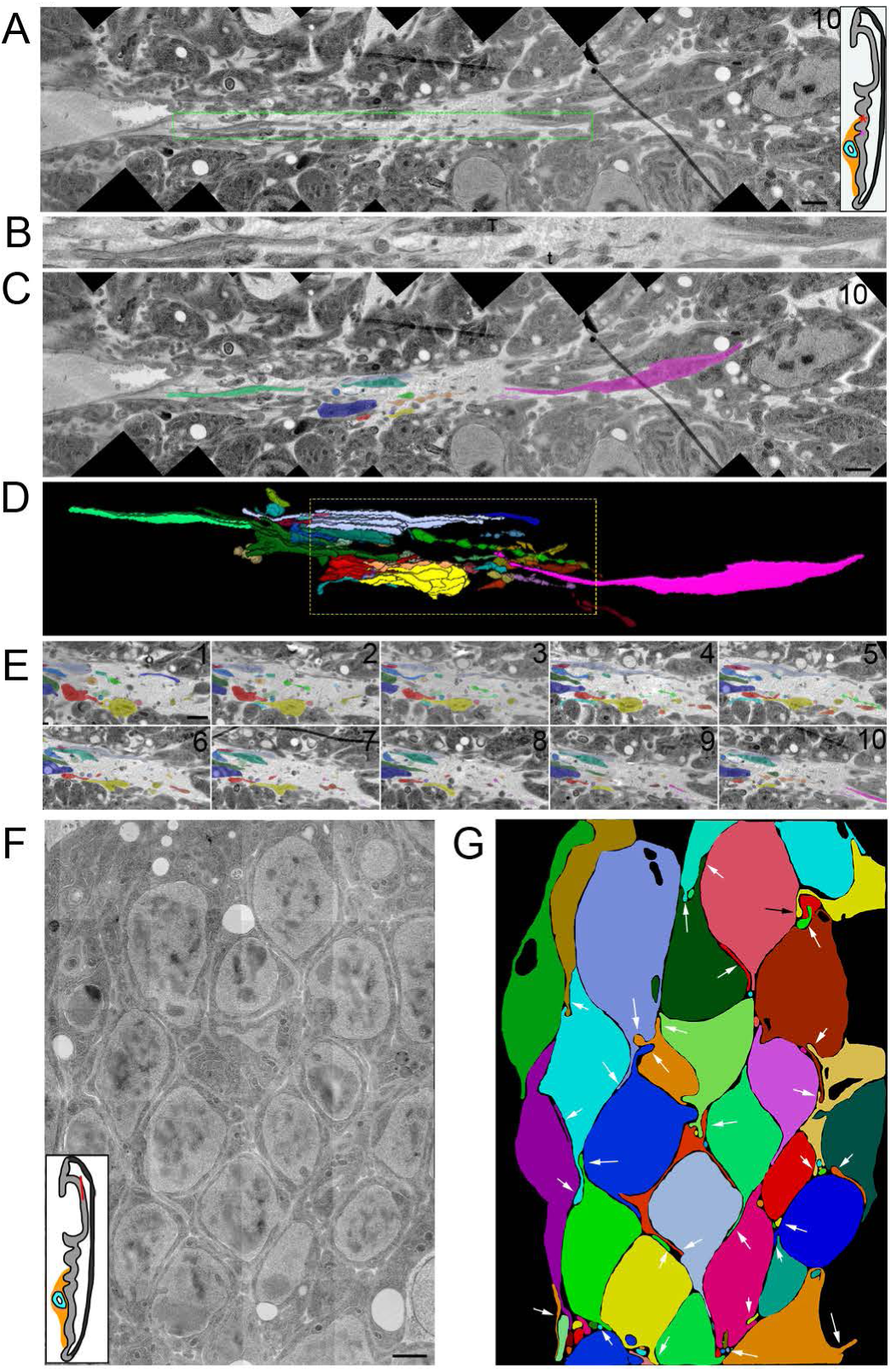
Cytonemes extend from basal and lateral surfaces of disc cells. **A)** One section of a ten section TEM series showing the basal side of the disc in a fold region (asterisk on disc diagram). Thick membrane extensions and thin cytonemes extend along the fold. **B)** An enlargement of the dashed green box in (A) with a mix of thick membrane extensions and thin cytonemes with constrictions and varicosities. **C)** Segmentation of individual membrane extensions and cytonemes. D) 3D-reconstruction of segmentation showing thick membrane extensions, thin cytonemes, and a thick membrane extension that narrows to a cytoneme (pink). **E)** Each of the 10 segmented sections used to produce the boxed region of D. **F)** Original and **G)** segmented image of one section of an 8 section TEM series showing cytonemes extending from lateral surfaces of disc cells (arrows). The red line on the disc diagram shows the area imaged. Scale bars: 1 µm.

Cytonemes also extend from basolateral sides of disc epithelial cells. In contrast to the cytonemes that extend from the ASP, myoblasts, and basal side of the disc, basolateral disc cytonemes are tightly confined by narrow extracellular spaces (Fig. 2F,G, Sup Fig 2). The paths of these cytonemes are defined by the contours of the cells and adjacent cytonemes.

### Cytonemes extend into invaginations of target cells

To investigate how cytonemes physically interact with target cells, we used SEM and TEM to track cytonemes projecting from their origins to cells they contact. 3D serial reconstruction of a SEM stack (Sup Fig. 3A) revealed a 13 µm long cytoneme the extends from a myoblast (green) past several myoblasts, and contacts a more distant myoblast (red) (Fig 3A). This cytoneme terminates in a cavity that envelops the cytoneme with target cell membrane (Fig. 3B). The diameter of the cytoneme shaft varies several times along the length of the cytoneme. These undulations are spaced at regular intervals in both the proximal regions far from the target cell and in distal regions that are inside the target cell cavity where the target cell membrane closely mirrors the shape of the undulating shaft. At the end of the cytoneme, however, there is a gap of as much as 82 nm between the cytoneme tip and the cell membrane (Fig. 3C).

**Figure 3.**
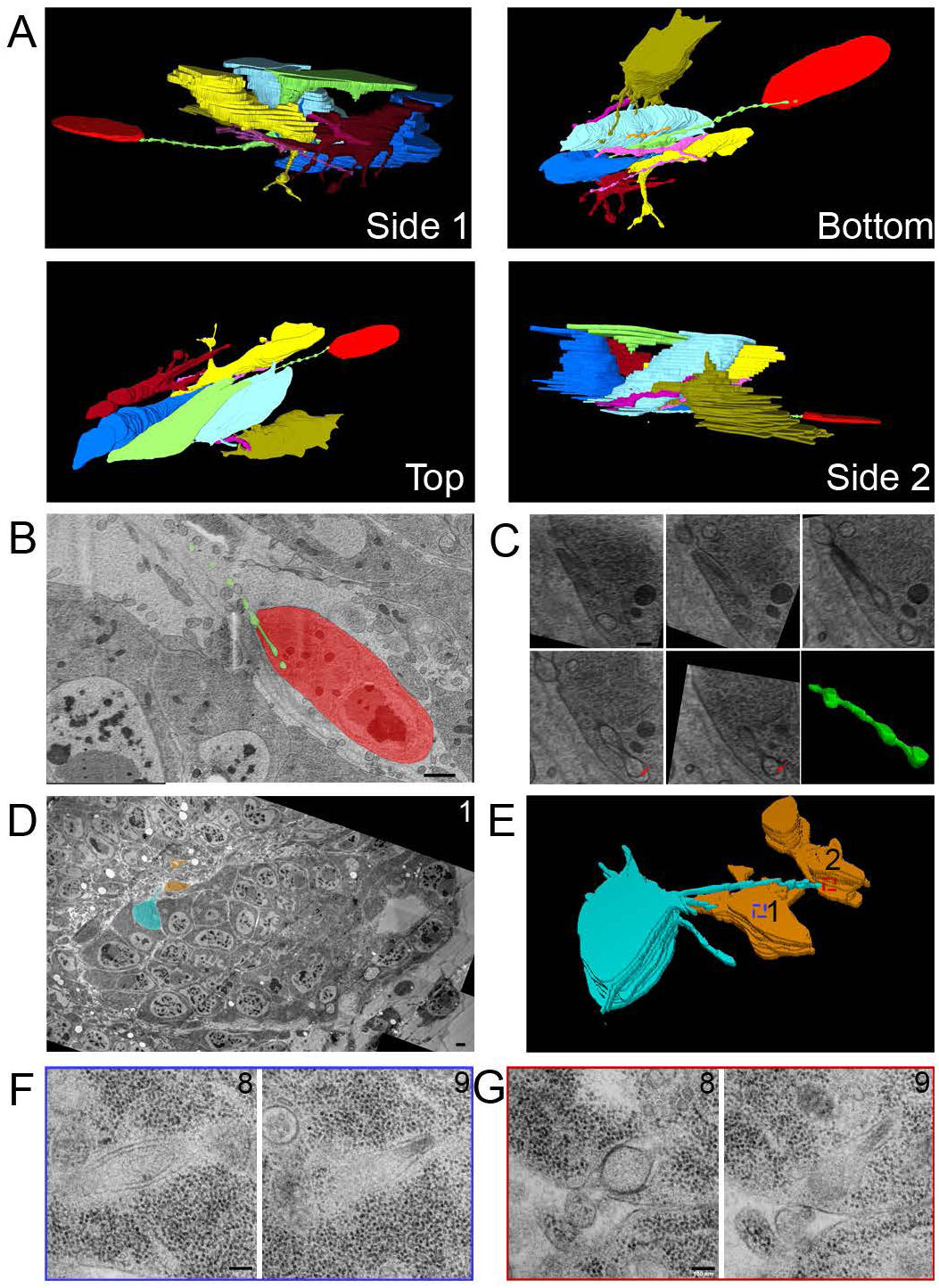
Myoblast and ASP cytonemes end in membrane invaginations of myoblasts. **A)** Four perspectives of a 3D reconstruction of myoblasts and their cytonemes from 61 SEM images spanning a depth of 6.4 µm. The cytoneme extended by the green cell end in an invagination of the red cell membrane. **B)** A segmented section showing the ending of the green cytoneme in a membrane invagination of the red cell. The white box and line on the disc diagram show the area imaged. **C)** Higher magnification SEM images of the five serial sections containing the portion of the green cytoneme inside the red cell invagination. The last panel contains a 3D reconstruction of the part of the cytoneme inside the invagination, showing that it retains the same shape and structure when connected to the target cell. **D)** One segmented TEM image highlighting an ASP cell (cyan) with two cytonemes which connect to two different myoblasts (orange). **E)** 3D reconstruction of 10 segmented serial TEM images showing ASP cytonemes in contact with and invaginating into two myoblasts (labeled 1 and 2). The blue and red boxes mark the XY position of the end of each cytoneme and the area imaged at higher magnification. High magnification images showing the last two sections of the cytonemes in contact with **F)** myoblast 1 and **G)** myoblast 2. Scale bars: 1 µm (B), 200 nm (C), 1 µm, 100 nm (F,G).

Images of ASP cells from serial TEM (Sup Fig. 3B, Fig. 3D) and 3D reconstruction (Fig. 3E) reveal similar structures. The ASP cell highlighted (cyan) in these panels extends cytonemes that project toward two myoblasts (orange). A higher magnification view of the two sections containing the most distal portions of these cytonemes has cytoneme termini surrounded at close proximity by myoblast membranes (Fig. 3F-G). An area of high density is present at the point of contact between the membrane of myoblast (#2) and the cytoneme (Fig. 3G, section 8). In contrast to the rounded ending of the burrowing cytoneme in Figure 3C, the ends of both cytonemes in this section are narrow (section 9) relative to the more proximal portions (section 8).

Previous work identified cytonemes connecting disc epithelial cells to each other (Huang and Kornberg, 2015; González-Méndez et al., 2020), and our SEM and TEM analysis also identified cytonemes that extend between ASP cells. Partial 3D reconstruction of four neighboring ASP cells from serial SEM images reveals cytonemes that extend from two cells (blue and cyan) which terminate within membrane invaginations of another (yellow). The blue cell cytoneme bifurcates into two branches within the yellow cell and both branches extend further into the yellow cell (Fig. 4B). The yellow cell extends a cytoneme that terminates within membrane invaginations of the purple cell (Fig. 4B-D).

**Figure 4.**
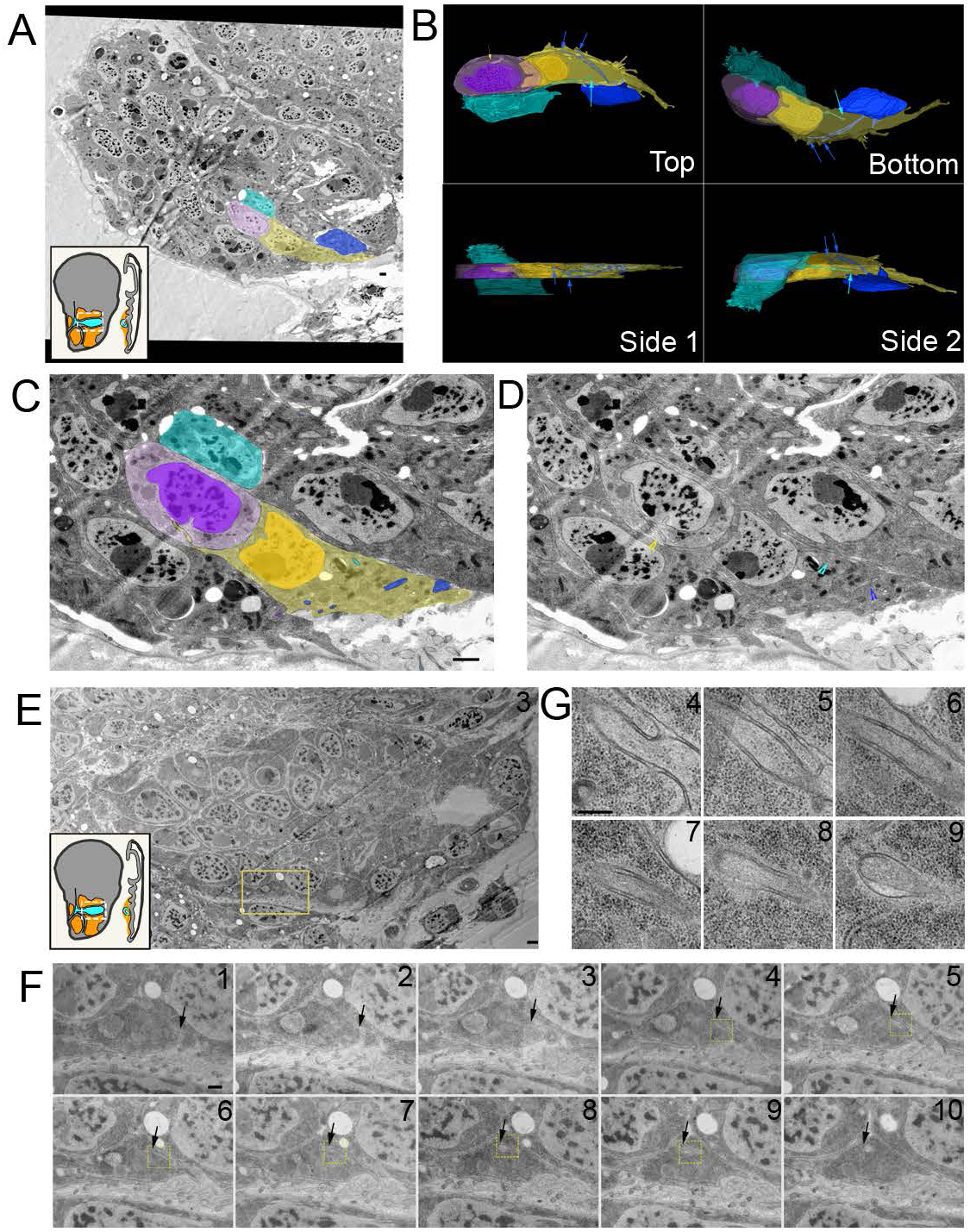
ASP cytonemes connect to neighboring ASP cells in membrane invaginations. **A)** Four neighboring ASP cells visualized by serial SEM were segmented. The cyan cell was segmented throughout the entire stack, while the other cells were partially segmented to allow for visualization of their cytoneme connections in 3D. **B)** Four perspectives of the 3D reconstructed ASP cells showing cytonemes from the blue and cyan cell ending within an invagination of the yellow cell, while the yellow cell projects a cytoneme ending within the purple cell. One **C)** segmented and **D)** non-segmented section showing cytonemes inside membrane invaginations of neighboring cells. Arrowheads point to the cytonemes of corresponding color. **E)** A TEM image of the ASP and surrounding myoblasts (location represented by white box and line shown on disc diagrams) shows a cytoneme from one ASP cell inside an invagination a neighboring cell. **F)** A series of 10 images (enlargements of the boxed region of E) show the cytoneme and its path (marked by arrows) within the membrane of the target cell. Boxed regions show the area imaged at higher magnification of sections 4-9 shown in **G**. Scale bars: 1 µm (A), 500 nm (C), 1 µm (E), 500 nm (F), 200 nm (G).

We also examined inter-ASP cell and inter-disc cytonemes by TEM. An example of an inter-ASP cytoneme and its connection to a neighboring cell is shown at high magnification in the series of micrographs in Figure 4E-F. This cytoneme-cell connection is similar to the connections of inter-myoblast cytonemes (Fig 3A-C) and ASP to myoblast cytonemes (Fig 3D-G). The cytoneme and cell membranes are closely apposed, except for a gap at the end of the cytoneme that is 45 nm wide at its largest point (Fig. 4G, section 9). Disc cytonemes, like ASP and myoblast cytonemes, also extend into membrane invaginations of target cells. Figure 6A shows a cytoneme originating at the lateral surface of a disc cell (cyan) and terminating within an invagination near the apical surface of a neighboring disc cell. This cytoneme bifurcates into two branches that both terminate within the target cell (Fig. 5B). A small uniform space separates the cytoneme and cell membrane throughout the length of their apposition. A cytoneme-cell connection with similar topology present near the basal side of the disc is shown in Figure 5C. This disc cell cytoneme terminates within a membrane invagination near the basal surface of its neighbor (Fig. 5C). In this instance, the gap between the cytoneme and cell membranes is also small and uniform.

**Figure 5.**
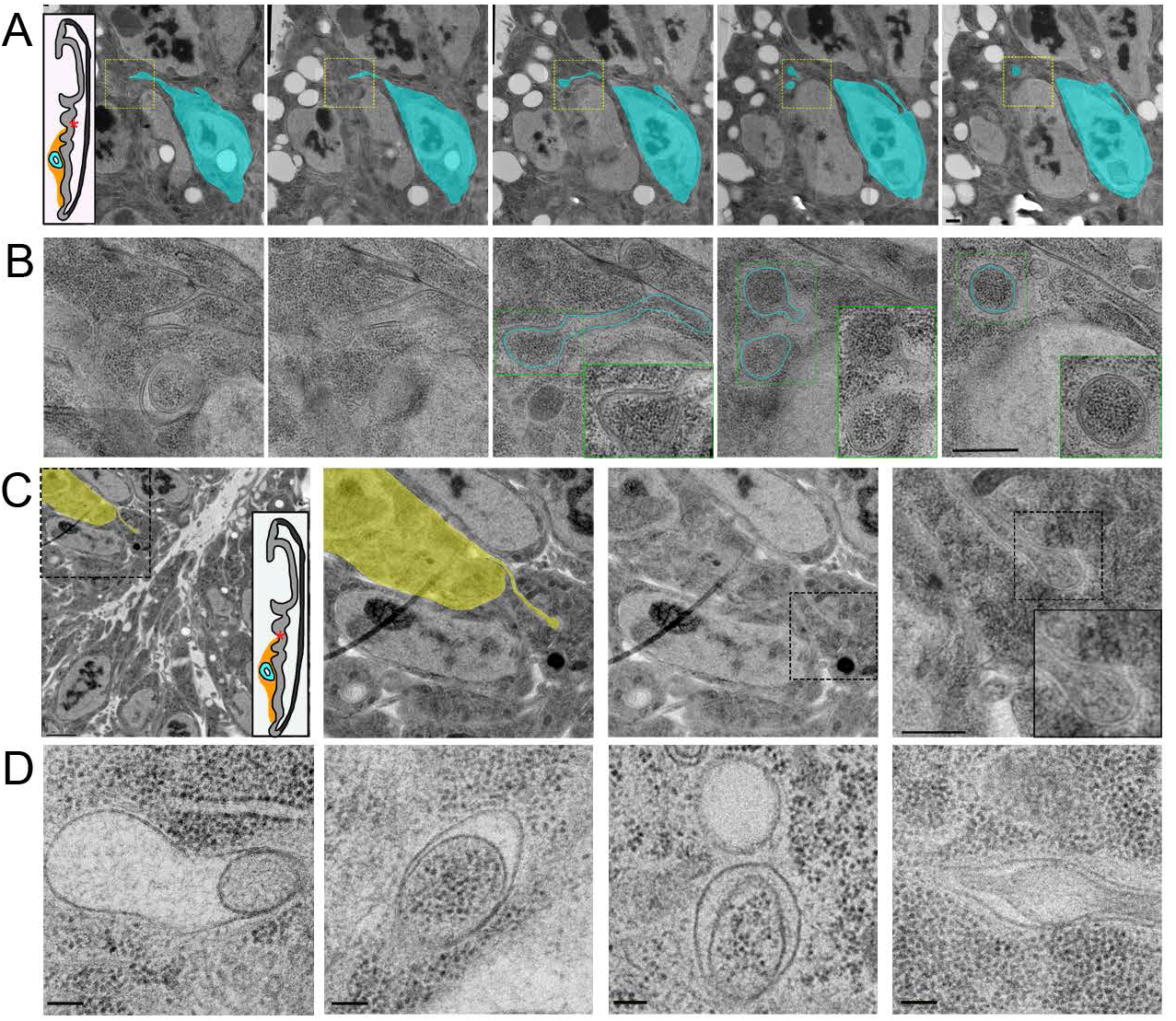
Disc cytonemes extend into invaginations near the apical and basal surfaces. **A)** A series of five TEM images showing a disc epithelial cell (cyan) projecting a cytoneme which ends in a membrane invagination of a neighboring disc cell. The region imaged is shown as an asterisk on the disc diagram. Part of the target cell’s apical surface and microvilli are seen in the upper left comer. The yellow box represents the area imaged at higher magnification shown in **B)** where the path of the cytoneme in the invagination is seen. Inside the invagination the cytoneme branches off into two terminal varicosities. **C)** A TEM image shows a disc epithelial cell (yellow) projecting a cytoneme from its lateral surface which ends in an invagination of the membrane of a neighboring disc cell near its basal surface. The area imaged is in the fold indicated on the disc diagram. Progressive enlargements of the boxed regions are shown from left to right. The middle two images are the same with and without segmentation of the cytoneme projecting cell. D) TEM images of four separate endings of cytonemes within invaginations of target cell membranes showing the variability in the size of the space between the end of the cytoneme and target cell membranes. Scale bars: 500 nm (A,B), 2 µm (C, panel 1), 500 nm (C, panel 4), 100 nm (D).

**Figure 6.**
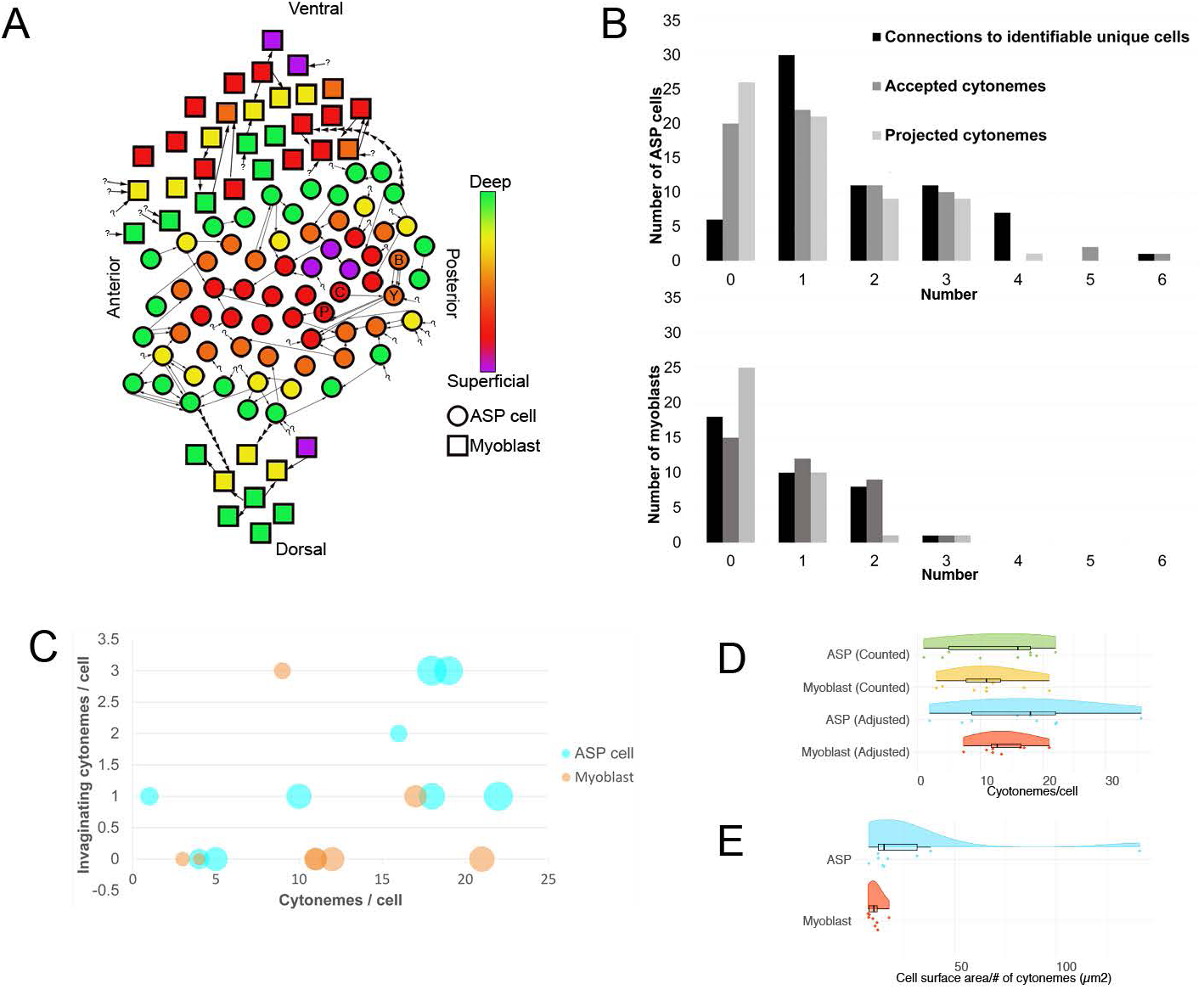
Intra-tissue cytonemes predominate in the ASP epithelium and myoblast population. **A)** A map of the approximate position of 66 ASP cells (circles) and 37 myoblasts (squares) contained within the serial SEM stack and their connections to other cells. The color indicates the z depth of each cell within the stack. Arrows begin at the cell from which a cytoneme originates and point towards the cell a cytoneme ends within. Question marks indicate invaginated cytoneme that could not be tracked back to a cell body. Positional labels (e.g. anterior) refer to the direction of the underlying disc compartments. The four cells segmented in Fig 4A-C are indicated on the map with letters corresponding to their segmentation color (B- blue, Y- yellow, C- cyan, P- purple). **B)** For each ASP cell (top) or myoblast (bottom), the number of connections (either sent or received invaginating cytonemes) to other verifiably unique cells, the number of invaginating cytonemes accepted, and the number of invaginating cytonemes projected to other cells was determined and graphed as a histogram. **C)** For nine 3D reconstructed ASP cells (cyan) and eight 3D reconstructed myoblasts (orange), the number of total cytonemes was compared to the number of those cytonemes that made contact and invaginated another cell. The area of the dot representing each cell scales with the surface area of the reconstructed cell. **D)** The number of cytonemes counted for each reconstructed cell, along with the number of expected cytonemes if all cells were contained fully within the stack are shown as individual data points and summarized as box plots with the probability density function. The adjustment factor is based on the surface area of each cell compared to the surface area of a completely reconstructed cell of the same type. **E)** The cell surface area per cytoneme for each reconstructed cell.

To obtain a more complete understanding of the prevalence and variations of inter-ASP cell and inter-myoblast cytonemes, the 66 ASP cells and 37 myoblasts visible in the SEM stack (Figs. 1,4, Supplemental Fig. 1) were examined and each cytoneme was tracked from its terminus inside an invagination back to its originating cell body. Not all cytonemes could be traced back to a source cell, either because they extended outside the imaged area or the images lacked sufficient resolution. A rough 2D map of the ASP and surrounding myoblasts was created by placing each cell in an approximate XY position and assigning a color corresponding to Z position within the stack (Fig. 6A). Four pairs of ASP cells and one pair of myoblasts reciprocally project cytonemes to each other. Five pairs of ASP cells and one pair of myoblasts connect by more than one cytoneme projected by one cell in the pair. Many cells both project and accept cytonemes, but some only project (14 ASP cells and 4 myoblasts) or accept (20 ASP cells and 14 myoblasts). Only three ASP cells project cytonemes which connect to myoblasts (Fig. 6A). By excluding connections made by accepted cytonemes that could not be traced back to a source cell and by counting repeated connections (multiple cytonemes between the same two cells) as one connection, the number of identifiable unique connections was counted for ASP cells and myoblasts (Fig. 6B). The cell with the most unique identifiable connections was an ASP cell linked to five other ASP cells and one myoblast. 60 of the 66 ASP cells connect by cytonemes to another identifiable cell; 19 of 37 myoblasts link to another identifiable cell, either by the presence of a cytoneme from another cell in an invagination or by extending one (Fig. 6B). ASP cells accept 90 cytonemes and myoblasts accept 33. ASP cells project 70 cytonemes that end in the invagination of another cell; myoblasts extend 15 (Fig. 6B). These results reveal that very few cells lack cytoneme-mediated cell-cell contacts, that inter-ASP cell cytoneme networks connect ASP cells, and that ASP cells that have large apposed membrane surfaces amplify the area of cell-cell contact with invaginations that juxtapose cytoneme and cell membranes.

The ASP and myoblast cell map shows only cytonemes that terminate in a target cell invagination. To estimate the percentage of total cytonemes this type represents, 9 typical ASP cells and 8 typical myoblasts were reconstructed in 3D, and the total number of cytonemes was compared to the number of invaginating cytonemes for each cell (Fig. 6C). The percentage of cytonemes which invaginate varies between cells (20.07% ±30.8 for ASP cells and 4.9% ±10.9 for myoblasts). Because most of the reconstructed cells have some portion outside of the imaged area, either in the XY or Z direction, the number of cytonemes counted for those cells is likely incomplete. To adjust for this, the surface area of a complete cell for each type was divided by the surface area of each incomplete cell and this factor was used to estimate the total number of cytonemes: 16.71 ±9.43 for ASP cells and 13.76 ± 3.95 for myoblasts (Fig 5D). Both the counted and adjusted number of cytonemes are shown in raincloud plots (Allen et al., 2019). The cell surface area that corresponded to projection of one cytoneme tended to be smaller for myoblasts than ASP cells (Fig. 6E), indicating cytonemes are more dense on myoblasts, even though the total number is similar to that of ASP cells.

Although the presence of cytonemes extending into invaginations of target cells is consistent for all three cell types, some aspects are variable. Whereas a space separates all the cytoneme:cell membrane juxtapositions, in some areas the space is better described as a gap (Fig, 6D, Sup Fig. 4). Both the size and shape of these gaps varies, but the differences are not characteristics of the particular cell type that either projects or receives a cytoneme.

### Periodic variations in shaft diameter create distinct varicosities and constrictions

Based on the general understanding of filopodia structure, we expected to find that cytonemes in our preparations would have shafts with a uniform and consistent diameter. Instead, we observed that with the exception of short (< 1 μm) extensions, all cytonemes emanating from ASP, myoblast, and disc cells have diameters that oscillate between narrow constrictions and well-defined wider sections (varicosities) (Fig. 7A,B). The constrictions have prominent actin filaments and little cytosol (Fig. 7C, green arrow), and the varicosities have more cytosol, including ribosomes (Fig. 7C, red arrow). No microtubules were identified in cytonemes, consistent with previous CM experiments. Although the presence of repeating sequences of constriction/varicosity/constriction/varicosity is a common feature of the cytonemes we observed, the shape, size, and spacing is variable (Fig. 7D,E and Supplemental Fig. 5). Cytoneme termini also vary, having widths that are as narrow as the constrictions or that have an expanded, bulbous structure, similar to varicosities. Bulbous termini are present in cytonemes that are entirely extracellular as well as cytonemes that burrow into membrane invaginations. The EM images do not distinguish differences in composition between the bulbous termini and shaft varicosities (Fig. 5D).

**Figure 7.**
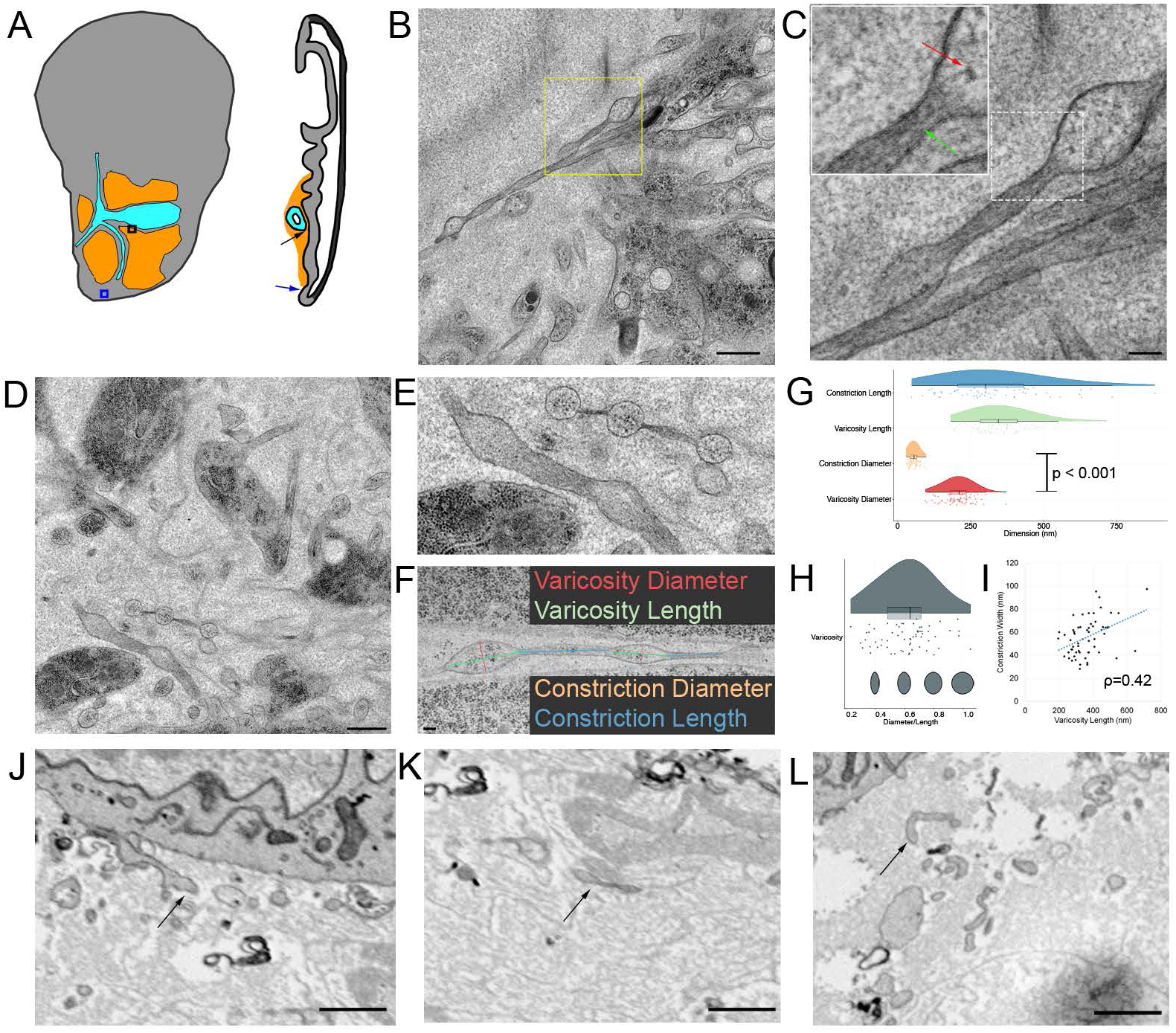
Cytoneme diameters vary between regions of varicosities and constrictions. **A)** Boxes and arrows indicate the imaged area shown in B (blue) and D (black). **B)** TEM image of two exemplar cytonemes with regions of varicosities and constrictions extended by disc epithelial cells (blue arrow in A). **C)** Higher magnification view of cytonemes corresponding to dashed box in (B). The inset is an enlargement of the area shown in the white dashed box. Ribosomes (red arrow) are visible in a varicosity, while actin filaments (green arrow) are visible in the constriction. **D)** TEM image of cytonemes among myoblasts (black arrow in A). **E)** Enlargement of two cytonemes from (D) showing the variance in size and shape of varicosities and constrictions. **F)** The width and length of 64 varicosities and 60 constrictions of cytonemes projected by the ASP and myoblasts were measured as shown. Box whisker plot of **G)** cytoneme dimensions and **H)** ratio between width and length of varicosities (dark grey) and constrictions (light grey). **I)** Plot of constriction width versus varicosity length showing a moderate linear correlation (p=0.42). **J)** SEM image of a chemically fixed cytonemes extending (arrow) from a disc epithelial cell with wide and narrow sections. **K)** SEM image of a portion of a chemically fixed cytoneme (arrow) with two varicosities separated by a constriction. **L)** SEM image of a chemically fixed cytoneme that does not show signs of having wide or narrow regions. Scale bars: 500 nm (B), 100 nm (C), 500 nm (D), 100 nm (F).

To quantify the shapes of constrictions and varicosities, we imaged cytonemes that project from ASP cells and myoblasts by serial SEM and analyzed 64 varicosities and 60 constrictions (Fig. 7F). Diameters were measured at the midpoint of each varicosity and constriction; because of their shape, the midpoint diameter of a varicosity is equivalent to the maximum diameter. Varicosity and constriction lengths are similar and both are varied (Fig. 7G). The period length (the length between midpoints of successive varicosities) was 716.3 ±206.6 nm. Varicosity diameters (206 ± 51.8 nm) are almost four times larger than constriction diameters (55.9 ±16.2 nm). The diameter to length ratio of varicosities is a measure of variability in varicosity shape (Fig. 7H). Most varicosities are ovoid, although some are more round (i.e. diameter/length equal to 1) and some more elongated. The relationship between varicosity length and constriction width is moderately positive (Fig. 7I), but no other relationship between varicosity or constriction width, length, or shape was found.

To determine whether chemical fixation also preserved the structure of the cytonemes in HPF- erived sections, we examined sections of chemically fixed wing discs. Chemical fixation followed by reduced osmium-thiocarbohydrazide-reduced osmium (ROTO) staining and serial SEM examination revealed cellular and organellar membranes with good preservation and demarcation (Supplemental Fig. 6). Although cytonemes and cytoneme membranes were also preserved (Fig. 7J; Supplemental Fig. 6), regions of varicosities and constrictions are not well-defined (compare Fig. 1F and Fig. 7J-L).

### Large membrane extensions contain cellular organelles

Although some cytonemes extend directly from cells as thin projections, many cells project wider membrane extensions that narrow down gradually into thin cytonemes (Fig. 8A, red arrow). Within these wide membrane extension regions, mitochondria, rough ER and membranous compartments are visible (Fig. 8A,B; Supplemental Fig. 7). We did not observe mitochondria or rough ER in the thinner cytoneme segments. This result is consistent with confocal images of cytonemes that extend from ASPs that overexpress either mito-GFP (and membrane bound mCherry; Fig. 8C) or RFP tagged with KDEL, an ER localization signal (and membrane bound GFP; Fig. 8D). Both the mitochondria and ER markers are prevalent in the wider regions of membrane extensions that are proximal to the cell body, but not in thinner regions of membrane extensions. ASP membrane extensions do not all have the same proportions of mitochondria and ER markers.

**Figure 8.**
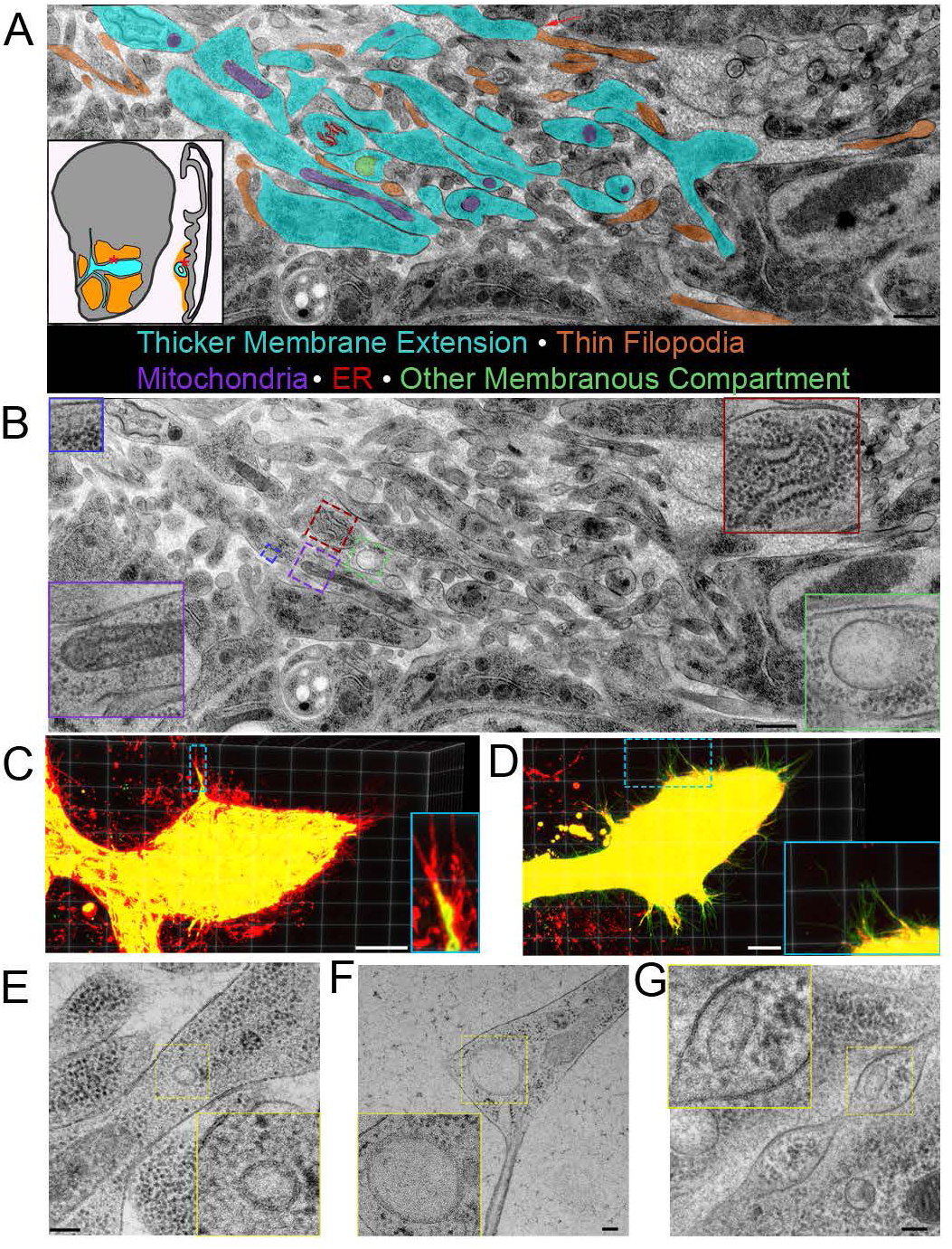
Thick membrane extensions contain mitochondria, ER, and membranous compartments. **A)** Partially segmented TEM image of thick membrane extensions (cyan) and thin cytonemes (orange) running between ASP and myoblasts (white box and line on disc diagrams). Some cells project thick membrane extensions that narrow into thin cytonemes (red arrow). Thick membrane extensions contain mitochondria (purple), rough ER (red), and other membranous compartments of different sizes (green). Membrane extensions and cytonemes that were in cross section were not segmented. **B)** TEM image shown in A without segmentation. Insets show enlarged examples of mitochondria (purple border), rough ER (red border), and other membranous compartments (green and blue border). 3D projection of confocal image stack of ASP expressing **C)** CD8-mcherry and Mito-GFP and **D)** CD8-GFP and KDEL-RFP. **E)** TEM image of E) a small membranous compartment inside a thick area of a membrane extension, **F)** a larger membranous compartment within a thick portion of a membrane extension and bifurcation into thin extensions, and **G)** a membranous compartment inside a varicosity of a cytoneme. Scale bars: 500 nm (A, B), 20 µm (C,D), 100 nm (E-G).

Round or oval membranous compartments are also present in many wide membrane extensions (Fig. 8E,F). These membranous compartments range widely in size from <50 nm to nearly 800 nm. Membranous compartments also varied in shape, with some being circular (Fig. 8E,F) and others ovoid (Fig. 7G). Membranous compartments in thin parts of the extensions are infrequent (Fig. 7G).

### Synaptic vesicles are present at immature neuron-myoblast synapses

Although membranous compartments similar in size to synaptic vesicles were infrequent in cytonemes or in the cytosol of the target cell surrounding cytoneme termini, we did observe synaptic vesicles concentrated at neuronal synapses in the sections of wing discs. Although there are no previous reports of synapsing neurons in third instar wing discs, we discovered an axon that synapses with multiple myoblasts. This axon was observed to stain with *α*-horseradish peroxidase (HRP) antibody in >40 discs (Fig. 9A,B), and was identified in one disc by TEM (Fig 9C-E). HRP staining highlights the axon which extends along the disc stalk and among the myoblasts, and branches at several points (Fig. 9A). Varicosities are present along the axon (Fig 9A). Immunostaining confirmed the presence of synaptotagmin within the neuron; staining was especially enriched at some varicosities (Fig. 9B).

**Figure 9.**
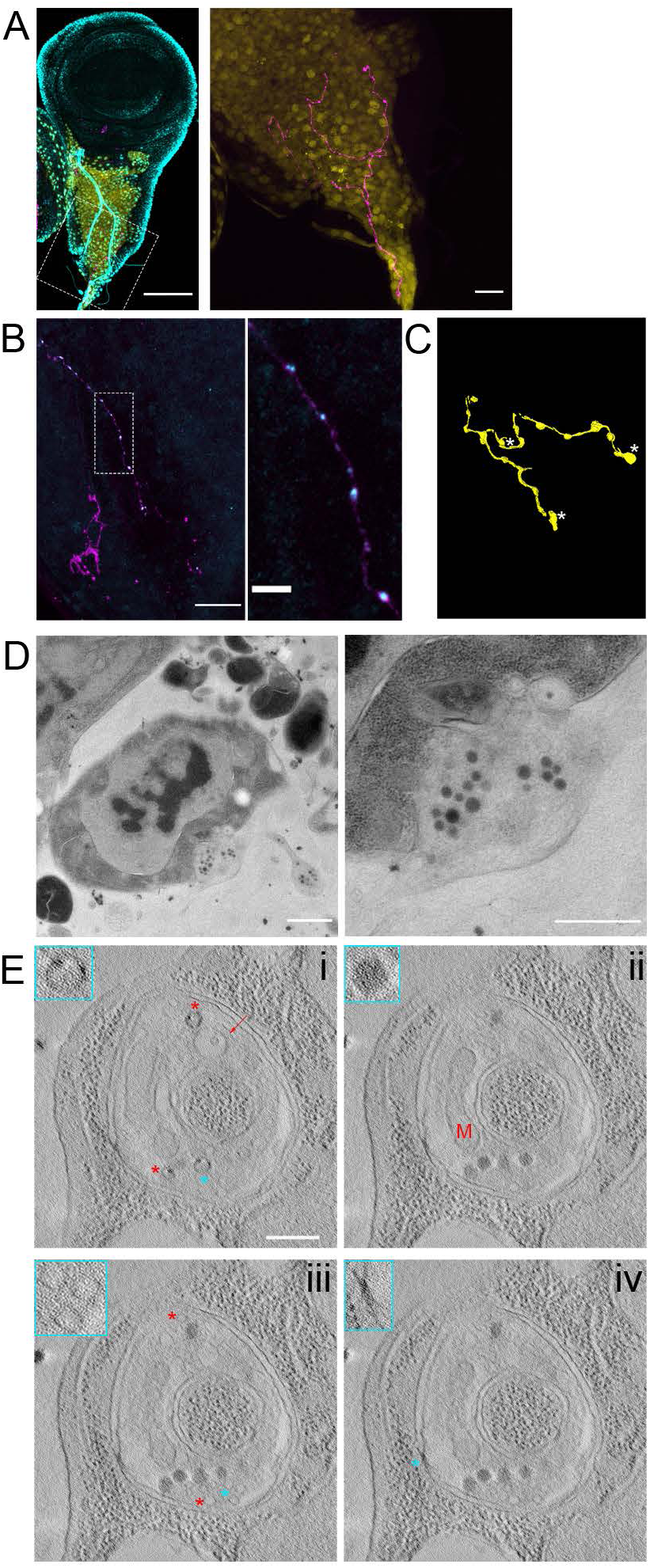
Synaptic vesicles are preserved in a neuron synapsing with wing disc myoblasts. **A)** A fixed wing and leg disc from an L3 larva expressing GFP (yellow) under the control of 1151-Gal4 labeled with DAPI (cyan) and HRP-AF647. The neuron (magenta) can be seen among the myoblast layer (yellow) of the wing disc. The right panel is shown without DAPI signal. **B)** A portion of a fixed disc stained for HRP (magenta) and synaptotagmin (cyan). Confocal images are shown as Z projections. **C)** A portion of the myoblast region was imaged by TEM in 40 serial 100 nm section. The montaged and aligned images were used to segment and 3D reconstruct the neuron. The location of four identified synapses along the neuron are indicated by an arrow, corresponding to the synapse shown in (D), or by asterisks. **D)** A TEM image showing a section of one of the observed neuron-myoblast synapses. At higher magnification (right), vesicles are apparent. **E)** Images from a tomographic tilt series containing the synapse shown in (D). Tomography revealed vesicles with electron dense cargo appearing as puncta on the inner edge of vesicles (i, asterisks), a membranous compartment containing a smaller vesicle (i, arrow), a mitochondrion (ii, M), dense core vesicles (ii, asterisk), clear synaptic vesicles (iii), and electron dense junctions (iv) between the neuron and myoblast membranes. The insets outlined in cyan show enlarged portions of the image that correspond to cyan asterisks, while red asterisks indicate other examples of the feature shown in the inset. Scale bars: 100 µm (A, left), 20 µm (A, right and B, right), 1 µm (D, left), 500 nm (D, right and E).

The myoblast region with the axon was analyzed in 40 100 nm sections by TEM, and images were used to segment and reconstruct a portion of the neuron in 3D (Fig. 9C). Four synapses between the neuron and myoblasts were identified (locations marked in Fig. 3C) and were imaged at higher magnification. Synaptic vesicles were found clustered at synapses (Fig. 9D), and along the length of the axon, especially at varicosities. A tomographic tilt series of one of the four synapses (Fig. 9D, Supplementary Fig. 8) revealed vesicles with different contents (Fig. 9E), including vesicles with cargo which appeared as dense puncta (Fig. 9E i, asterisks), vesicles contained within a larger membranous compartment (Fig. 9E i, arrow), dense core vesicles (Fig. 9E ii), and clear vesicles (Fig. 9E iii). A ∼50 nm cleft separates the neuronal and myoblast membranes except at electron dense junctions (Fig. 9E iv). Mitochondria (Fig. 9E ii) were also observed at synapses. The synapsing portion of the neuron is a spheroid varicosity positioned within a concave pocket of the myoblast membrane (Supplementary Fig. 8). A portion of the myoblast membrane is enveloped in an invagination of the neuron membrane (Supplementary Fig. 8, arrows). The features of these synapses suggest that they are not fully mature, as they lack the characteristic T-bar structure of mature Drosophila synapses. The presence of vesicles at neuronal but not cytoneme synapses in our samples, shows that vesicles are preserved but that cytoneme synapses do not have the same type of standing reservoir of vesicles. This suggests that vesicles at cytoneme synapses are much fewer and/or transient than at neuronal synapses.

## Discussion

Imaging filopodia has historically been limited by technical impediments. The small diameter of filopodia, which has been approximated at ∼200 nm, limits detection by standard transmitted light microscopy, and because their optical properties are similar to cell bodies, they are essentially invisible where they extend over cells. They are also preserved poorly by chemical fixatives. Recent technical innovations, however, now make it possible to see and investigate filopodia in ways that have not previously been possible. Contemporary confocal microscopes have better spatial resolution and more sensitive detectors, filopodia can be brightly marked with fluorescent proteins, and there are efficient methods for mosaic expression of fluorescent, genetically-encoded proteins. With these tools, light-emitting filopodia can be detected where they extend over non-fluorescent cells.

The Drosophila wing disc system has many attributes that make it favorable for viewing cytonemes. Its three most abundant cell types – disc epithelia, trachea epithelia and mesenchymal myoblasts - have low auto-fluorescence. It is relatively flat and has few cell layers. It can be isolated and mounted quickly without fixation for microscope viewing such that cytonemes extending across epithelia lie in a small number of optical planes. In addition, there are many ways to make genetic mosaics, and signaling systems that communicate within and between the tissues are well characterized. Studies carried out during the past twenty years have revealed many novel features of cytonemes in the wing disc system.

Cytonemes are abundant in the wing disc, and many are specialized for signaling with components such as receptors, transporters and channels. As dedicated signaling organelles, they are designated cytonemes. And as imaging and detection methods have improved since cytonemes were discovered, the inventory of number and variety of cytonemes has increased substantially. Cytonemes have been detected extending from every cell type in the disc system. They are dynamic, vary in length to >100 μm, make synaptic contacts that are essential for exchanging signaling proteins and for signaling, and are regulated such that their numbers correlate with signaling level. Nevertheless, although much has been learned, the limitations of fluorescence microscopy leave many questions unresolved. We do not know, for instance, how many cytonemes a particular cell makes, the precise architecture of the actin-based extensions, the macromolecular contents, or the fine structure of the connections they make with target cells. To address these questions, this study imaged cytonemes and cytoneme connections using HPF, SEM, and TEM to better resolve the ultrastructure and contents of these specialized filopodia. Our findings not only provide novel insights into these essential signaling organelles, but in addition have important general implications for the nature of filopodia and the process of synaptogenesis.

By capturing cells with HPF and analyzing cell morphologies at nanometer resolution, we determined the inventory of cytonemes which extend from individual epithelial and mesenchymal cells at an instant in time. Every cell we analyzed has at least one cytoneme, with ASP cells and myoblasts averaging 17 and 14, respectively. The cell with the highest number is an ASP cell with 22. ASP cytonemes are exclusively basal and basolateral and have varied orientation. The basal surface of most ASP cells projects at least one thick membrane extension that transitions to a thin cytoneme that tracks distally along the basal surface of its distal neighbor cell. Myoblasts lack apparent apical/basal polarity, but have an expanded, oblong medial region from which both thick projections and cytonemes extend. All prominent projections extend radially in the plane of the disc surface and without apparent polarization with respect to anteroposterior or dorsoventral disc axes or the proximodistal ASP axis. In this landscape of epithelia and mesenchyme, we conclude that projections extend the reach of every cell.

Fluorescence microscopy identified cytonemes that extend between anterior compartment disc cells and posterior compartment disc cells (Chen et al., 2017; González-Méndez et al., 2017), between ASP and disc cells (Sato and Kornberg, 2002; Roy et al., 2011), and between myoblasts and ASP cells (Huang and Kornberg, 2015). However, cytonemes connecting cells of the same type in the myoblast layer or ASP had not been previously investigated. Our EM analysis reveals that whereas most cytonemes do not contact another cell, when they do it is most commonly with cells of the same tissue.

Based on our previous studies and studies of others (Huang et al., 2019; Junyent et al., 2020), we expected that connections between cytonemes and target cells might share structural features with neuronal synapses. Neuronal synapses are characterized by specializations of the pre- and postsynaptic compartments, including the postsynaptic density (Kennedy, 2000), a cleft between the two membranes containing electron dense complexes (Zuber et al., 2005), and clusters of vesicles in presynaptic active zone (Harris and Weinberg, 2012). The presence and precise characteristics of these specializations vary with the stage and size of the synapse. During synaptogenesis in insects, the first contact between synaptic partners is initiated by growth cone filopodia that burrow into their target (Bastiani and Goodman, 1984), and in vertebrate immature synapses, electron densities and vesicle-rich active zones are absent (Stelzner et al., 1973; Altman, 1971; Bunge et al., 1967; Glees and Sheppard, 1964; Fröhlich and Meinertzhagen, 1982). The cytoneme connections we imaged burrow into target cell invaginations and do not have the electron densities and vesicles that are characteristic of mature neuronal synapses. There are regions of high density between the cytoneme and target cell membranes (Fig. 3G and 4I) that are similar to specializations that have been observed in developing house fly brains (Fröhlich and Meinertzhagen, 1982). These cytoneme connections appear to share features with the early, maturing neuronal synapses which acquire structural components stepwise (Fröhlich and Meinertzhagen, 1982). The lack of a standing pool of vesicles at cytoneme synapses does not invalidate evidence that vesicles are important for cytoneme signaling (Huang et al., 2019; González-Méndez et al., 2020); rather it suggests that cytoneme vesicles are few and transient.

Structures with cardinal features of neuronal synapses were identified in our EM preparations. Although synapsing neurons have been identified in the wing disc system during the pupal but not larval stage (Jan et al., 1985), the use of collagenase prior to fixation in this previous study may have resulted in the loss of the neuron we identified that contacts myoblasts in 3^rd^ instar discs. In addition, this neuron is thin, and the increased brightness of fluorophores and increased sensitivity of detectors available today may also explain why it had not yet been detected. The soma of this neuron is outside of the wing disc, but its axon appears to extend into the hypodermal stalk of the wing disc and through the myoblast layer. The connections between the neuron and myoblast have synaptic vesicles, mitochondria, and electron dense junctions, that are consistent with synaptic function, although they lack the T bars that are characteristic of mature Drosophila synapses.

The cytoneme-cell connections we detected penetrate several microns into cell invaginations, a feature shared with signaling cellular projections such as those extended of granulosa cells of the mouse ovarian follicle (Baena and Terasaki, 2019) and growth cone filopodia of the grasshopper embryo (Bastiani and Goodman, 1984). Membrane invagination by close-ended TNTs has also been observed (Sartori-Rupp et al., 2019)(Sartori-Rupp et al., 2019), and invaginating structures are common at mammalian synapses (Petralia et al., 2015, 2018). Even sponges elongate processes that make invaginating contacts with other cells, although they do not have neurons (Petralia et al., 2015).

Penetrating cytonemes are mirrored closely by the cell membrane except at the end where a gap is frequent. This gap ranges in size and shape, perhaps representing different stages of maturing cytoneme-cell connections. Growth cone filopodia that selectively invaginate the membranes of other growth cones and neuron cell bodies have a space along both the sides and tip of the filopodia that increases during maturation (Bastiani and Goodman, 1984). It is possible that the cytonemes we observed with large amounts of space around the terminus represent a more advanced stage of cytoneme-cell interaction.

In both extracellular space and invaginations, cytoneme shafts have distinct regions of wide diameter varicosities and narrow diameter constrictions. Although filopodia have been examined by EM before, to our knowledge, this is the first study to examine filopodia in high pressure frozen tissue by serial EM. High pressure frozen filopodia have been visualized in cross section before, but the shape along the length of filopodia was not assessed. (Lak et al., 2015). Chemical fixation during EM processing causes shrinkage of tissues and distortion of cellular and organellar membranes. This may explain why filopodia imaged after chemical fixation have more uniform diameter and less smooth membranes. The microvilli that we detected in our samples are mostly several microns long, similar to short cytonemes, and have a diameter range between 50-250, comparable to cytonemes, but every microvillus has a uniform diameter, lacking varicosities or constrictions. These observations indicate that the non-uniform diameter of cytonemes is not an artifact of our methods.

The varicosity/constriction cytoneme architecture has features reminiscent of en passant synapses, which have variably spaced varicosities at synaptic sites (Giachello et al., 2012; Shepherd et al., 2002). TEM analysis of human intraretinal ganglion axons has identified varicosities with very similar shapes as cytoneme varicosities, although the axon diameter is >10x that of cytonemes (Wang et al., 2003). And although neuronal varicosities store synaptic vesicles and other synaptic machinery, most cytoneme varicosities have high ribosomal content and few contain vesicles or membranous compartments. Bulges have been observed in TNTs by SEM (Scholkmann et al., 2018), but their contents, including vesicles and other membranous compartments, distinguish them from cytoneme varicosities.

Although some cytonemes extend directly from the cell, others extend from thicker membrane projections where mitochondria, rough ER, and membrane bound compartments are common. These organelles are extremely rare in cytonemes. Ribosomes, in contrast, are abundant and widely disseminated in cytonemes, and are often at the same density as the cellular cytosol. This suggests that thick membrane projections and filopodia are both cytosolic extensions of the cell, with any organellar exclusion based on the ability of a membrane extension to accommodate the size of an organelle rather the selective filtering. This is contrasts with primary cilia, which control content but have no ribosomes.

## Materials and Methods

### High Pressure Freezing (HPF)

Prior to dissection, planchets (Ted Pella, Redding, CA) were coated with 1-hexadecane (Sigma) and thoroughly dried. Wing imaginal discs were dissected in either PBS or Grace’s Insect Media (Thermo Fisher Scientific) from wandering third instar larva. Using forceps, the bottom third of the larva was removed. One tine of a forceps was inserted into the mouth, and the larva was turned inside out along the tine using the other forceps. Fat bodies, intestines, and dorsal trunks of the trachea were removed carefully. The wing disc was pulled away from the carcass and transferred via capillary action between forceps to a drop of PBS or media on a staging dish. 20% Bovine Serum Albumin solution made with either PBS or Grace’s Insect Media was used as cryoprotectant. A 0.1 mm “A” planchet was partially filled with BSA solution. Approximately 100 ul of BSA solution was added to the PBS or media drop containing the disc. The disc was transferred serially to two drops of BSA solution before being transferred to the planchet. The level of BSA solution in the planchet was adjusted to a slight convex meniscus with forceps. The planchet was transferred to the specimen holder and the flat side of “B” planchet was placed on top of the specimen containing planchet. The sample holder was transferred to the Baltec HPF machine and frozen. This process was performed as quickly as possible in order to reduce the time from dissection to freezing and exposure of the disc to BSA solution.

### Freeze Substitution

Freeze substitution was done according to the Super Quick Freeze Substitution protocol. After high pressure freezing, planchets containing samples were placed in cryotubes containing acetone-based media with 4% OsO_4_, 0.1% uranyl acetate, 2% MeOH, and 5% H_2_O. No more than three samples were placed in any one tube. The media had been pre-frozen and stored in liquid nitrogen. At all times after freezing and prior to freeze substitution (FS), samples were kept at liquid nitrogen (LN_2_) temperature by performing transfers in LN_2_ or LN_2_ vapor. Prior to FS, a metal heater block with slots for 13 mm tubes was cooled with LN_2_ in a rectangular foam ice bucket. Cryotubes containing the samples and FS media were transferred into one of the end rows of the LN_2_-filled block. The temperature was monitored by wrapping a temperature probe around a tube filled with acetone which was then placed in the block. To begin FS, the block was turned on its side within the ice bucket so that the row with the tubes was closest to the bottom of the bucket, and so that the tube caps were facing one wall of the ice bucket. The LN_2_ was poured out of the ice bucket and the uncovered bucket was placed on a shaker and agitated at 100 rpm. The bucket was shaken until the temperature of the tubes reached 2-4° C. The FS process took on average 2 hours.

### Resin Infiltration

Samples were washed five times with glass distilled acetone (Electron Microscopy Sciences (EMS), Hatfield, PA). During the last wash, the tubes were placed on a rocker for 5 minutes. The samples were gradually infiltrated with Epon/Araldite (EA) resin through progressive steps (25%, 50%, 75%) of EA/acetone mixture. Epon/Araldite resin consisted of 6.2 g Eponate 12, 4.4 g Araldite 502, 12.2 g DDSA. 1 ml of EA/acetone mixture was pipetted into the tube after removal of the previous step. Uncapped tubes were placed in a loud cooled microwave under vacuum (Ted Pella) and were microwaved with 150mW for 1 minute. For 25% and 50% steps, tubes were rotated for 1 hour at room temperature, after which they were briefly centrifuged to ensure the samples were at the bottom of the tube. Some samples came out of the planchets on their own during rotation, the ones that did not were visualized using a dissecting scope and removed using a needle tip to carefully trace around the border between the sample and planchet and gently lift. 75% EA/acetone was added to the samples, which were then microwaved and rotated overnight. The next day, the resin mixture was replaced with 100% E/A and rotated for 1 hour. The samples were placed in a new tube containing accelerated E/A (same recipe as above with 0.8 ml BDMA added) and rotated for 3 hours with exchanges of fresh E/A every hour. Between each exchange, samples were briefly centrifuged down. Samples were then flat-embedded between two Teflon-coated slides, with Parafilm used as a spacer, and baked at 60° C for 48 hours.

### Chemical Fixation

#### Section Preparation

For TEM, a Reichert-Jung Ultracut E microtome was used to cut 100 nm thick sections with a diamond knife (Ultra 45°, Diatome, Hatfield, PA) which were collected onto fomvar film coated Copper/Rhodium slot grids. Sections were post stained with 2% uranyl acetate and Reynold’s lead citrate.

For SEM, 60 nm sections were cut using a diamond knife (Ultra 45°, Diatome) and collected using an automated tape-collecting ultramicrotome. The tape was transferred to silicon wafers and carbon coated (Baena et al., 2019).

#### Electron Microscopy

Transmission electron microscopy was performed using a Tecnai 12 microscope (FEI, Hillsboro, OR) operating at 12 kV. Images were acquired with Gatan Microscopy Suite (GMS3) software and a Gatan U895 4k x 4k camera. Montage imaging was performed using SerialEM.

Tomography was performed on a Tecnai 20 microscope (FEI, Hillsboro, OR). IMOD software was used to reconstruct the tilt series.

SEM imaging was performed using two microscopes: a Zeiss Sigma field-emission scanning electron microscope in backscatter mode at 8 ekV(Fig 1 and 6), and a FEI Verios field emission scanning electron microscope in backscatter mode at 5 ekV (Fig 3).

The TrakEM2 module of FIJI ImageJ was used to montage images, align serial sections, segment images, and produce 3D reconstructions of segmentation. Images acquired on the Zeiss SE microscope were pre-aligned using the Linear Stack Alignment with SIFT ImageJ plugin before being imported into the TrakEM2 module.

#### ASP Connection Map

ASP cells within the 172 image serial SEM stack were assigned a number to track their connections. Each cell was examined to identify all cytonemes which invaginated with its membrane, and when possible the cytonemes was tracked back to the originating cell. The rough XY positioning of each cell was mapped and each cell was assigned to one of five Z depth colors determined by the placement of its nucleus within the stack. Green cells had the majority of their nuclei within sections 1-40 (deepest sections), yellow cell nuclei correspond to sections 41-80, orange cell nuclei correspond to sections 81-120, red cell nuclei correspond to sections 121-160, and purple cell nuclei correspond to sections 161-172. Arrows mark the direction of the connection only (originating at the filopodia sending cell and pointing toward the cytoneme receiving cell) and do not indicate information about the length or path of the cytoneme.

#### Confocal Microscopy

Live imaging: Wing imaginal discs were dissected as described above in Grace’s Insect Media. Discs were mounted by placing the disc in a drop of media on a coverslip with the ASP side down. Media was slowly pipetted from the drop until the level was such that the surface tension had just begun to compress the disc against the coverslip. The coverslip was inverted onto a concavity slide with small amounts of silicone vacuum grease applied to the area surrounding the slide depression. ASP were imaged using 488 and 560 wavelength lasers, an inverted laser scanning confocal microscope (Olympus FV-3000), a 40X Oil Objective, and GaAsp PMT detectors. Image stacks were 3D rendered with the ImageJ plugin ClearVolume.

Immunohistochemistry: The posterior third of the larva was removed, the cuticle was inverted, and fat bodies were removed. Carcasses were fixed in 4% paraformaldehyde for 25 minutes. After 3 washes of PBS, 0.2% Triton-X in PBS (TBST) was used to permeabilize cells for 10 minutes followed by blocking with Roche blocking reagent diluted 1:5 in TBST for 30 minutes. Antibodies were diluted in blocking reagent and incubated overnight at 4°C. After three 10 minute TBST washes, secondary antibody anti-mouse AlexaFluor555 (1:500) (Invitrogen), or Alexa405 conjugated phalloidin (1:400) were diluted in blocking reagent and incubated with the carcasses for 45 minutes at room temperature. After three 10 minute TBST washes, the wing discs were detached from the trachea and gently separated from the carcass. Two strips of double-sided tape were used to provide space between the slide and coverslip. Discs were mounted in Vectashield mounting reagent (Vector Labs). Confocal images are shown as maximum Z projections of multiple confocal slices.

Primary antibodies: Alexa647 conjugated anti-horseradish peroxidase (1:100) (Jackson Immunoresearch) and anti-synaptotagmin (1:5) (Developmental Studies Hybridoma Bank).

### Fly Lines

Confocal microscopy: *btl*-Gal4 (Roy et al., 2014), 1151-Gal4 from K VijayRaghavan (Roy and VijayRaghavan, 1997), (UAS-mcd8:GFP, UAS-mcd8:mCherry, UAS-RFP:KDEL (Bloomington #30910), UAS-mito:GFP (Bloomington #8443), UAS-GFP/Tm6B. Electron microscopy: w^1118^.

### Statistical Analysis

Constriction and varicosity widths were compared with a Student’s t test.

## Supplemental Material

Supplemental figures show serial section images of images found in Fig. 1-5 and Fig. 7-9. Supplemental Video 1 shows 360° rotations of the 3D reconstructed cells shown in Fig 1D and E.

## Acknowledgments

We thank Danielle Jorgens, Reena Zalpuri, and Kent McDonald at the University of California Berkeley Electron Microscope Laboratory for training and advice in HPF and TEM techniques. We thank Richard Fetter for advice, sharing of custom software, and help with tomography data acquisition.

The authors declare no competing financial interests.

## Author Contributions

**Brent Wood:** Conceptualization, Formal Analysis, Funding Acquisition, Investigation, Project Administration, Visualization, Writing-original draft. **Valentina Baena**: Investigation, Writing-review and editing. **Hai Huang**: Investigation, Writing-review and editing. **Mark Terasaki**: Investigation, Resources, Writing-review and editing. **Thomas Kornberg**: Conceptualization, Funding Acquisition, Resources, Supervision, Writing-review and editing.

